# Variation in developmental rates is not linked to environmental unpredictability in annual killifishes

**DOI:** 10.1101/2020.03.16.993329

**Authors:** P. K. Rowiński, W. Sowersby, J. Näslund, S. Eckerström-Liedholm, K. Gotthard, B. Rogell

## Abstract

Comparative evidence suggests that adaptive plasticity may evolve as a response to predictable environmental variation. However, less attention has been placed on unpredictable environmental variation, which is considered to affect evolutionary trajectories by increasing phenotypic variation (or bet-hedging). Here, we examine the occurrence of bet-hedging in egg developmental rates in seven species of annual killifish, which originate from a gradient of variation in precipitation rates, under three treatment incubation temperatures (21°C, 23°C, and 25°C). In the wild, these species survive regular and seasonal habitat desiccation, as dormant eggs buried in the soil. At the onset of the rainy season, embryos must be sufficiently developed in order to hatch and complete their life-cycle. We found substantial differences among species in both the mean and variation of egg development rates, as well as species-specific plastic responses to incubation temperature. Yet, there was no clear relationship between variation in egg development time and variation in precipitation rate (environmental predictability). The exact cause of these differences therefore remains enigmatic, possibly depending on differences in other natural environmental conditions in addition to precipitation predictability. Hence, if species-specific variances are adaptive, the relationship between development and variation in precipitation is complex, and does not diverge in accordance with simple linear relationships.

## INTRODUCTION

Organisms can cope with fluctuations in environmental conditions by means of phenotypic plasticity or bet-hedging (Crean and Marshall 2009; Simons 2011; Furness et al. 2015a). However, studies that investigate both plasticity and bet-hedging at the same time have largely been theoretical (e.g. Cooper and Kaplan 1982; DeWitt and Langerhans 2004; Marshall and Uller 2007; Scheiner and Holt 2012; Tufto 2015), and it is only more recently that empirical studies have begun to focus on validating these theoretical predictions (i.e. Bradford and Roff 1993; Richter-Boix et al. 2006; Marshall et al. 2008; Simons 2014; Furness et al. 2015a; Shama 2015). Understanding the evolution of these two adaptive strategies has become critical in the face of global climate change, which is changing not only environments, but also the predictability of environmental conditions (Robeson 2002; Thornton et al. 2014; Berg and Hall 2015).

For plasticity and bet-hedging to evolve, there must be variation in the fitness of individuals under different environmental conditions (i.e. phenotypes having differing environment optima; Simons 2011). Phenotypic plasticity is an environment-dependent trait expression, and given genetic variation in reaction norms (i.e. genotypes differ in their associated phenotypes depending on environmental conditions; Simons 2011), adaptive phenotypic plasticity is considered to evolve as a response to predictable environmental changes, for which there are reliable cues (Ghalambor et al. 2007). However, phenotypic variation *per se* can be adaptive, as over a range of environmental conditions, by chance alone, the phenotypes of some individuals may be close to the environment dependent fitness optima (i.e. bet-hedging) (Crean and Marshall 2009). Therefore, when environmental conditions are either not predictable, or non-transducible into developmental regulators, production of highly variable phenotypes will spread the risks associated with unsuitability to particular environmental conditions. Under this scenario, bet-hedging is considered to be an adaptive strategy (Clauss and Venable 2000; Crean and Marshall 2009).

The seemingly random nature of bet-hedging helps to ensure the survival of at least some offspring, which also results in reduced among-generation variation in reproductive success (Crean and Marshall 2009). Bet-hedging is considered to be a costly strategy, as there will be constant selection against a non-trivial part of the population that is not suited to current environmental conditions (Kussell & Leibler 2005; Beaumont et al. 2009). Further, while bet-hedging can theoretically evolve in any trait, it is hypothesized to be particularly relevant in traits related to juvenile establishment, in highly fecund organisms that do not engage in parental care, and also in organisms that inhabit areas prone to rapid environmental fluctuations (e.g. “*r*-selected” species, following MacArthur and Wilson 1967).

One textbook example of bet-hedging is observed in annual desert plants, which produce seeds with highly variable germination times, in response to unpredictable rates of precipitation (Venable 2007). In this case, if there is a drought in any given year, at least a subset of seeds will still likely germinate in the following years, and ensure survival of the lineage (e.g. Phillipi 1993; Evans et al. 2006; Venable 2007). Similarly, in animals, bet-hedging is observed in relation to important upcoming environmental events, such as the arrival of winter or the next rainy season, which may be difficult to predict. For example, differences in diapause length during harsh periods (e.g. winter or dry seasons) has been reported within many animal taxa, such as insects, crustaceans, rotifers, and in fish (Cáceres and Tessier 2004; García-Roger et al. 2014; Furness et al. 2015a; Hopper 2018). In these examples, not only variation in the length of diapause, but also variation in the proportion of individuals (often at the egg/embryo stage) that enter a diapause phase or not has been observed (Cáceres and Tessier 2004; García-Roger et al. 2014; Furness et al. 2015a; Hopper 2018).

Bet-hedging is therefore an adaptive increase in phenotypic variation, and under a bet-hedging scenario, trait variation is expected to correlate to environmental variation (Crean and Marshall 2009). Species often inhabit different environmental conditions that differ in both variability and predictability. Thus, by comparing different species or populations, which inhabit environments that vary in the predictability of particular environmental conditions or events, we can test for the generality of bet-hedging strategies (van Kleunen et al. 2014; Hopper 2018). For example, in plants, comparative studies have repeatedly demonstrated bet-hedging in seed dormancy (e.g. Philippi 1993; Evans et al. 2006; Venable 2007). Moreover, evidence from several studies suggests that differences in bethedging responses occur among animal populations and species (Marshall et al. 2008; Krug 2009; Nevoux et al. 2010; García-Roger et al. 2014; Polačik et al 2018). Yet, questions remain regarding the adaptive nature of these patterns, as few studies have assessed differences across species or populations that occur over a gradient of environmental predictability (but see García-Roger et al. 2014; Polačik et al. 2018).

Here, we investigate mean and variation in the developmental time of eggs in seven species of annual killifishes (Cyprinodontiformes, Aplocheiloidei). These species inhabit ephemeral freshwater bodies in Africa and South-/Central America, where both within- and between-season conditions are often highly unpredictable (Inglima et al. 1981; Genade et al. 2005; Furness 2016). Across killifishes, several species have independently evolved these specific adaptations to survive in ephemeral habitats. For instance, in these annual clades, eggs stay dormant (i.e. in diapause) buried in the substrate of dried-out pools until the next wet season (Furness et al. 2015b). However, even within an annual species, the duration of egg diapause is variable and not always obligatory, meaning that one clutch of eggs may consist of both directly developing and diapausing eggs (of varying duration) (Wourms 1972b). This contrast in development time has been suggested to constitute a bet-hedging strategy that maximizes fitness by spreading the risks associated with variability in the hydrological dynamics of ephemeral pools (Wourms 1972b; Polačik et al. 2014; Furness et al. 2015a). Interestingly, variation in the duration of killifish diapause has been found to be both under maternal control and governed by plasticity during embryonic development (Podrabsky et al. 2010; Polačik et al. 2016a; Pri-Tal et al. 2011; Furness et al. 2015a).

Using killifishes, we measured the duration of egg development in a standardized laboratory setting. Furthermore, we correlated species-specific means and variances of egg development time, with measurements of variation in precipitation rate, from the species native ranges during the onset of the rainy season. Eggs were reared under three different temperatures to assess the extent to which development time variation was stable across thermal conditions. The seven killifish species included in this study are representative of five of the annual clades that independently transitioned from a non-seasonal to a seasonal life-history (as indicated by presence of type II diapause).

clades that have evolved the annual life cycle. These species were also chosen because they originate from a gradient of environmental conditions, with clear differences in the predictability of precipitation rates. We predicted that species that have evolved under conditions which are relatively more unpredictable will have larger variation in egg development rates, compared to species that have evolved in areas with more predictable environments.

## MATERIALS AND METHODS

### Animal models

We used seven annual killifish species, *Gnatholebias zonatu*s (Myers 1935), *Millerichthys robustu*s (Miller and Hubbs 1974), *Nematolebias whitei* (Myers 1942), *Nothobranchius guentheri* (Pfeffer 1893), *No. kadleci* Reichard 2010, *Pituna schindleri* Costa 2007, and *Simpsonichthys constanciae* (Myers 1942), collected from 12 locations/populations (see Table S1, supplementary information 1 for details), and which are representative of five of the major independent evolutionary transitions between a non-annual and annual life cycle (Furness et al. 2015b; Gonzalez-Voyer et al. unpublished data). The experimental procedures were approved by the Ethical Committee in Stockholm, Sweden (license N132/15).

### Rearing of parental generation

The parental generation was hatched from eggs sourced from dedicated hobbyists, or from our own laboratory housed breeding groups. The parental generation were hatched under standardized conditions, and raised individually in 0.75 liter plastic containers with ramshorn snails (Planorbidae) to consume uneaten food, and java moss (*Taxiphyllum barbieri*) for shelter. They were all fed (3 times/day-weekdays, and 1 time/day weekends) with freshly hatched *Artemia salina* nauplii and reared under standardized conditions (ambient temperature 28°C, average water temperature 24.2°C ± 0.65 s.d). Tap-water (KH=4, GH=7, pH=7.5), with the addition of Jbl Biotopol (JBL GmbH & Co, Nauhofen, Germany) water conditioner was used to fill aquaria. The parental fish were pooled into groups of 3-4 individuals at one week of age, in one 0.75-liter plastic box. When they then reached 1cm in total length, they were transferred to 13-liter tanks and fed with a mixture of de-frosted Chironomid larvae and live *Artemia salina* nauplii. The 13-liter tanks were furnished with gravel, an empty terracotta flowerpot and a yarn mop. Water quality was maintained with an air-driven sponge filter, and twice weekly 80% water changes in the 0.75-liter plastic containers and weekly 25% water changes in the larger 13-liter tank.

### Breeding procedures

When females were noticeably mature, that is, with a rounded egg-filled belly, they were paired with a randomly chosen male and placed together in a 13-liter spawning tank for a period of 2-3 months (Information on the number of families, breeding males and females per species, and number of surviving eggs per family may be found in Table S2, supplementary information 2). Female annual killifish either bury or disperse eggs over a substrate, therefore a 0.75-liter plastic container, filled with either glass beads (Sargenta AB, Malmö, Sweden) or coco peat (Exo Terra, Hamm, Germany) was added into each breeding tank to provide substrate for egg laying. We found that glass beads facilitated more efficient egg retrieval, but these were not readily accepted by all species as an appropriate breeding substrate, so those species (n=3) where supplied exclusively with boxes filled with coco peat. In most species (n=6), some males were particularly aggressive, and were therefore grouped together with 2-4 females to dilute aggression among a higher number of females. Females were regularly switched into different tanks with a different male, residing in each for 2-3 months. Eggs were gathered weekly, either by sieving the glass beads through a net, or by laying the coco peat on a white plastic board and inspecting thoroughly for the eggs.

### Experimental treatment of the offspring generation

We included different temperature treatments during egg incubation to asses if developmental times are congruent across different thermal regimes. This is in line with Furness et al. (2015a), who investigated bet-hedging and developmental plasticity in *No. furzeri*, and found that percentage of eggs entering diapause differed depending on rearing temperature. However, while Furness et al. (2015a) investigated a desert species, and used rather high rearing temperatures, we used a a range of temperatures that likely is within the range of what all species would encounter in the wild. In this regard, fertilized eggs from each male tank (per female or female group), were partitioned into three different egg incubators (Lucky Reptile Herp Nursery II, Import Export Peter Hoch GmbH, Waldkirch, Germany) set to 21°C±0.15 s.d, 23°C±0.44 s.d, and 25°C± 0.33 s.d. All adult/parental fish were kept under a 12:12 light regime, and while the developing embryos were also kept in the same laboratory, they were housed in incubators and therefore not exposed to direct light. There are several recognized methods for incubating killifish eggs (see Polačik et al. 2016b), but to maximize the standardization of rearing conditions and to avoid artificially increased within-species variation, we chose to incubate eggs submerged in a Yamamoto solution (17 mM NaCl, 2.7 mM KCl, 2.5 mM CaCl2, pH set to 7 with NaHCO3; Valenzano et al. 2009 after Rembold et al. 2006; also successfully used by Furness et al. 2015a, b, 2016). In addition, 2 drops of 6.25 mM methylene blue and 5.33 mM acriflavine solution were added per 1 liter of Yamamoto solution, to prevent the occurrence of fungus and bacterial infections. For incubation, each embryo was transferred to a separate well containing the incubation medium, on a 24-well tray (TC-Plate Standard F, Sarstedt AG & Co. KG, Nümbrecht, Germany). The Yamamoto solution was changed twice a week to ensure clean and stable solution conditions.

### Data collection

Killifishes can have up to three distinct embryonic diapause stages, referred to as Type I (developmental arrest during early development, dispersed cell phase), Type II (38 somites present, beginning of organ development), and Type III (embryo developed and able to hatch) (Wourms 1972a; Furness 2015b). During Type II diapause, embryos are particularly resistant to dehydration stress, and this stage is only observed in annual killifish (Podrabsky et al. 2015; Furness 2015b; Furness 2016). Moreover, as Type II diapause allows for survival of harsh environmental conditions and may last for many months, the time until the end of Type II diapause is considered an important time-point for survival during the dry season (Podrabsky et al. 2015, Furness 2016). In annual killifishes, the eyes of embryos become pigmented after the Type II diapause phase (Wourms 1972a). Therefore, we visually inspected embryos weekly with a magnifying glass for the appearance of pigmented eyes. The period between the egg laying date and the appearance of pigmented eyes was then used as a proxy for developmental time. In total we collected 2567 eggs, out of which 24% (*N* = 614) survived until the eye-pigmentation stage. The majority of mortality occurred shortly after egg collection, with half of the mortality occurring during the first week of incubation (see Table S2 in supplementary information 2 for more information on the eggs that survived), presumably due to being unfertilized or inflicted with minor damage during collection, which may have exposed the embryos to oomycete (Saprolegniaceae) infections. At the end of experimental period (which ran between July 2017 and June 2019), 21 embryos did not show eye development. These undeveloped embryos, which were evenly distributed across the temperature treatments, belonged to two species, *G. zonatus* and *P. schindleri*, and were excluded from the analysis. We consider these exclusions very unlikely to have influenced our results, as they represented less than 1% of the total embryos included in our study, and around 5% of embryos collected for each of these two species.

### Estimation of precipitation variability

To assess the influence of precipitation variability on both embryo development time and variation in development time, we identified the exact coordinates of the collection localities of our lab populations from the killi-data.org archive (Huber et al. 2016), and retrieved site-specific climate data for each locality from the Local Climate Estimator software (New_LocClim, average length of time-series is 50 years; FAO 2018). The following variables were collected: *i*) mean among-year precipitation for the three first months of the rainy season, and *ii*) the among-year standard deviations in precipitation. We chose these specific parameters, as precipitation predictability during the rainy season should be key to complete pond-filling, and hence, crucial for embryo survival. We reasoned that high variation in precipitation rates during the early rainy season may result in ponds only partially filling. Partial filling may be enough to induce killifish eggs to hatch, but possibly not provide enough time to complete their entire life cycle, as a partially filled pool might desiccate quickly in times of drought. For each species, the first month of a rainy season was assumed to be the month when precipitation increased, following the dry season. The dry season was considered as a within year period of low precipitation, with average precipitation falling under 60 mm ∙ month^−1^ (Peel et al. 2017), during at least one month. In the case of *Ne. whitei*, the differentiation between dry and rainy season was not as obvious as other species, and indicated the existence of two separated rainy seasons. Hence, we averaged precipitation data for these two periods (Table S1, supplementary information 1). When our lab populations originated from more than one collection locality, we averaged the climate data for these populations (Table S1, supplementary information 1). The averaged climate data is therefore representative for the species, as different populations of the same species used in this study typically inhabit similar environments (Table S1, supplementary information 1). Annual killifishes may disperse across areas during flooding events, promoting gene flow between populations which would diminish differences in local adaptation between populations (Berois et al. 2015).

As variance scales to the mean, and as there are large divergences among the populations in the mean rainfall, we calculated species-specific coefficients of variation (*CV*) for monthly precipitation (average length of data series being 50 years), as the standard deviations multiplied by 100, divided by the mean, and averaged the monthly *CV*s for the first three months of the rainy season.

### Statistical models

To examine sources of variation in egg development time, we first fit two separate intercept-only models. The first model was fit on a subset of the whole data, when female identity was known (i.e. data from breeding tanks containing only one female, model 1a, Table S3, supplementary information 3), and included species, male, and female, as random effects. The second model was fit to the full dataset, including those observations where female identity was unknown (breeding tanks with both single females and female groups, model 1b, Table S3, supplementary information 3), and included species, male, and female/female group, as random effects. Of these models, the former model gave us an opportunity to estimate female effects, while the later model provided better estimates of species effects, given the larger dataset. We then calculated medians and 95% confidence intervals of male, female/female group, within-, and among-species posterior distributions of variation.

In order to *i*) correlate environmental predictability with means and variances of development time, and *ii*) compare among-temperature means and variances of development time, we first ran four intercept-only models, one model per temperature treatment, and one with pooled temperatures. All models included species as a random effect and allowed for separate residual variance among species (models 2-5, Table S3, supplementary information 3). From these models, we obtained full posterior distributions for species-, and temperature-specific means and variances of development time, which allowed for calculation of posterior distributions for *CV*s in developmental times. The posterior distributions for the *CV*s were calculated as the square root of posterior distribution for variances, multiplied by 100, divided by the posterior distribution for the mean.

In order to assess the significance of species-specific responses to temperature, we calculated the differences among the three temperature-specific posterior distributions for each species (Fig. 2 and 4). Significance was determined as a lack of overlap with 0.

Finally, to test if variation in development time increased with increasing variation in precipitation, as would be expected under a bet-hedging scenario, we modelled *CV*s of development time as a response variable, and a vector of precipitations *CV*s as an explanatory variable. This was done for the full posterior distribution of CV (*N*_iterations_ = 1000), to account for uncertainties in the estimates. In addition, for each iteration, we bootstrapped over species, in order to account for uncertainty associated with the included species (where, for example *M. robustus* could be classified as an outlier). This approach yielded posterior distributions of the slope of the regression, where significance was determined as a lack of overlap with 0. We used the same approach to examine if means of development time were associated with precipitations *CV*s.

### Potentially confounding factors

Maternal age has previously been found to affect egg development time in killifishes (Podrabsky et al. 2010; Pri-Tal et al. 2011; Polačik et al. 2016a). Therefore, we wanted to exclude the sources of maternal effects that could be viewed as experimental artefacts. Specifically, as pairing was preformed over an extended period (see Table S2 in supplementary information 2 for more information on the crossings), the age of parental fish differed across different full-sib families, which previously has been found to influence embryo development time (Podrabsky et al. 2010). Moreover, while our intention was to create full-sib offspring families, some males were more aggressive and multiple females were required in the mating tanks, which could potentially influence egg developmental trajectories. Hence, we initially ran a model with female age (continuous variable) and female group rearing (factor with two levels, single and group) as fixed effects. Species, male, and the female/female group identity, were added as random effects, in order to access if female age or group/single female breeding could constitute a source of maternal effects, and influence the results. However, as these factors were non-significant, they were not included in any of our inferential models.

### Model evaluation

Data was analyzed using Bayesian mixed-effect models (MCMCglmm-package for R; Hadfield 2010), in R version 3.4.4 (R Development Core Team 2015). In all models, we used flat priors for the fixed effects and locally non-informative priors for the random effects, at least 1∙10^6^ iterations, a burn-in of 5∙10^4^, and a thinning interval of 1000, which resulted in effective sampling size of 1000.

We diagnosed posterior distributions and model convergence by running three parallel chains using the Gelman–Rubin convergence criterion; the upper 97.5 quantile of the Gelman-Rubin test statistic was below 1.2 in all cases. All autocorrelations were within the interval −0.1 and 0.1.

## RESULTS

We found that most of the variation in development time was structured within species, as the residual variation accounted for 70% (95% CI: 36, 85) of the total variation (averaged across species). Among species variation accounted for 28% (95% CI: 12, 63) of the total variation, while genetic and maternal influences were negligible (male <1%, female < 1%, and female/female group <1%).

### Species- and temperature-specific length of development time

Development time differed significantly among species (Figure 1), and among different rearing temperatures within species (Figure 2), as indicated by non-overlapping 95% CI’s of the posterior distributions. However, there was no clear linear relationship between development time and precipitation *CV* (*β* = −0.0048; 95% CI: −0.82, 1.65; Figure 1). Furthermore, there was no general relationship between development time and rearing temperature, although in a subset of species, *S. constanciae, P. schindleri,* and *Ne. whitei,* higher temperature correlated with shorter development times (Figure 2).

**Figure 1.**
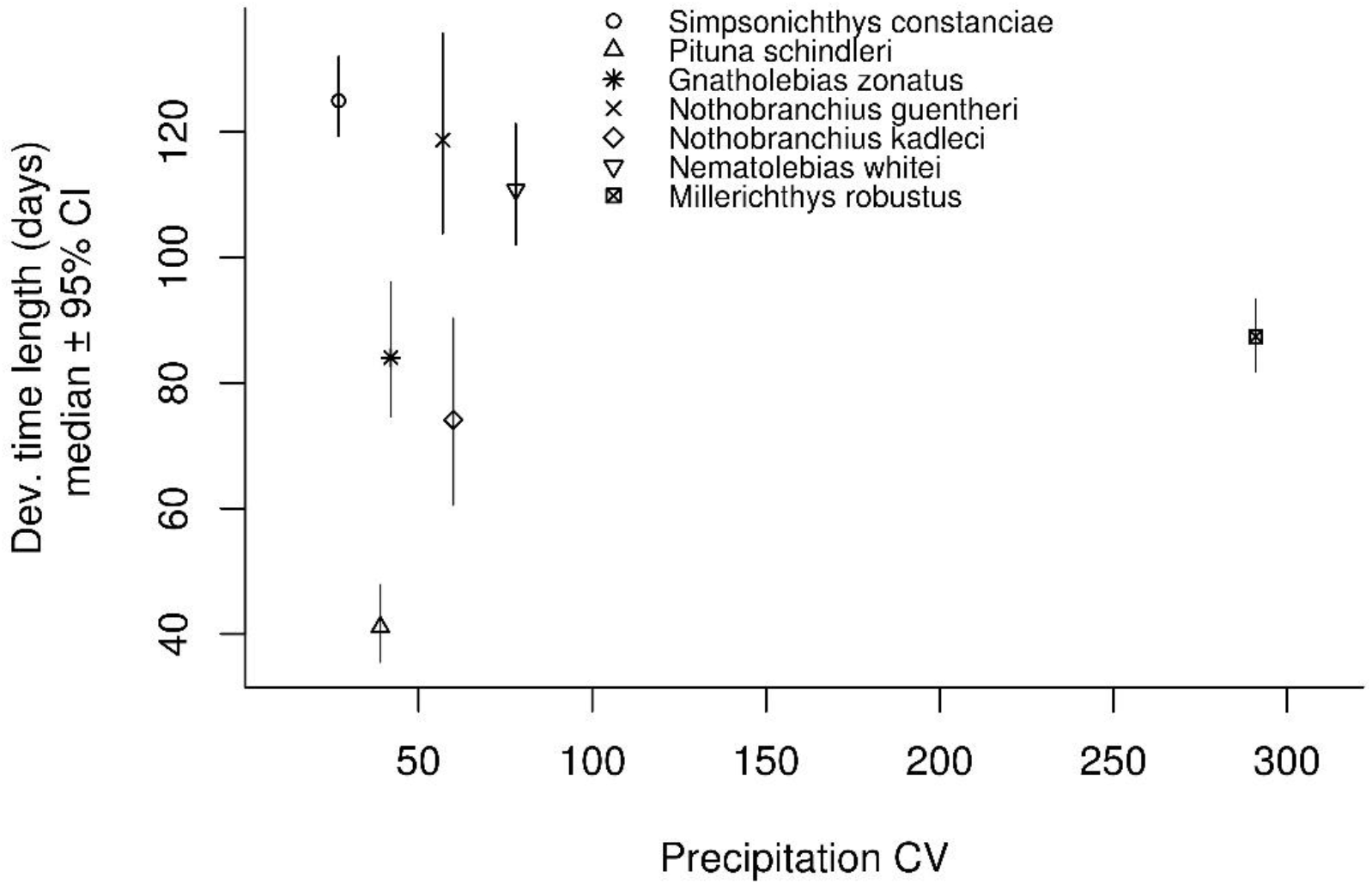
Species-specific medians of posterior distributions of development time length, and their 95% credibility intervals (y-axis), against precipitation CV (x-axis).

**Figure 2.**
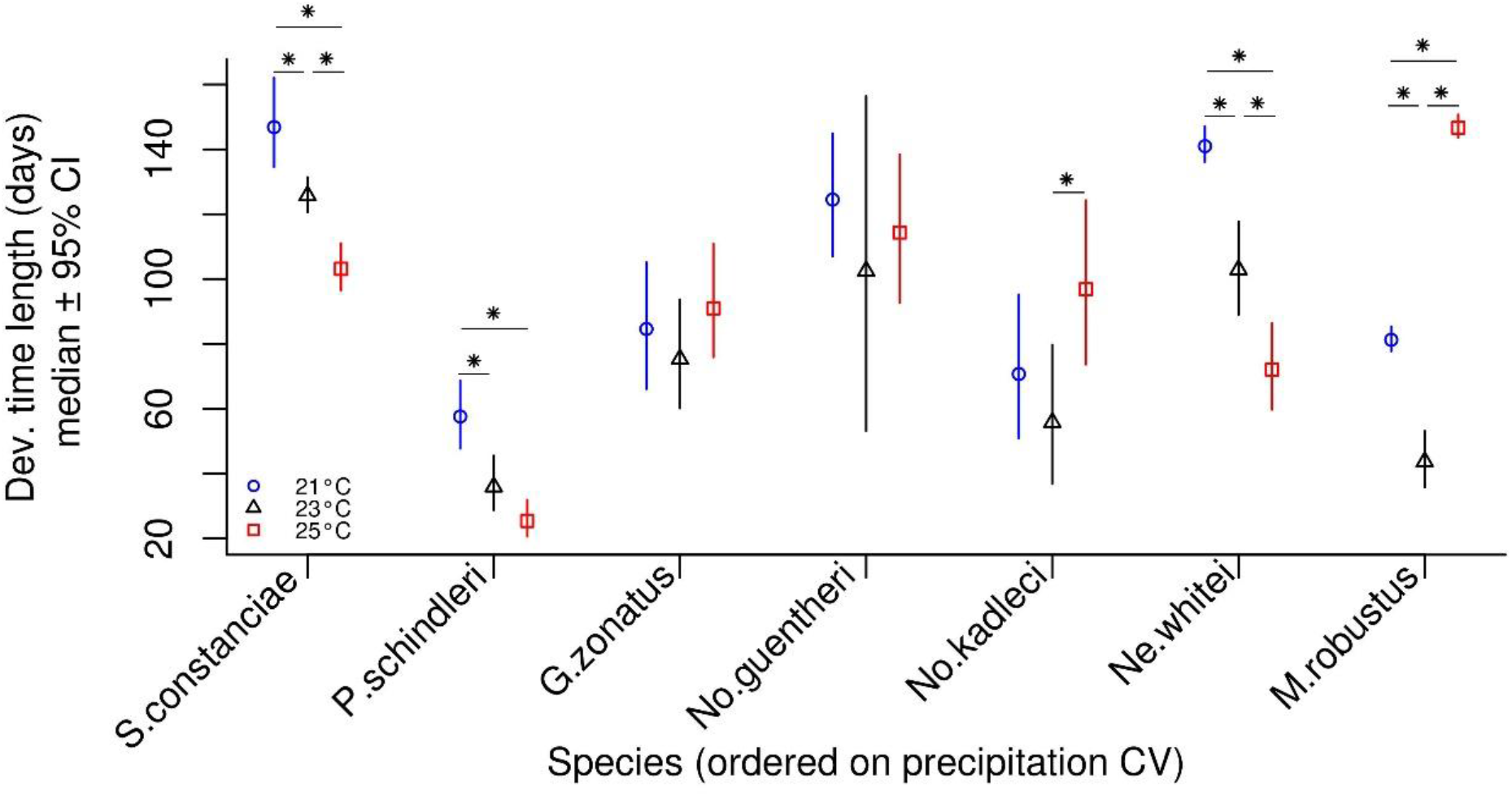
Species- and rearing temperature-specific medians of posterior distributions of development time length, and their 95% credibility intervals (y-axis). Species are ordered on a categorical x-axis scale, according to precipitation *CV* values, from the lowest (left) to the highest (right), for clarity of the results. Star symbols indicate significant within-species differences between the temperature treatment groups. A similar plot, but with continuous x-axis scale can be found in the supplementary material 4, Fig. S1.

### Species- and temperature-specific variation in development time

Development time *CV*s differed significantly among species (Figure 3), and among different rearing temperatures within species (Figure 4, *S. constanciae, Ne. whitei, M. robustus*), as indicated by nonoverlapping 95% CI’s of the posterior distributions. There was no clear linear relationship between development time *CV* and precipitation *CV* (*β* = 0.077; 95% CI: −0.56, 1.13; Figure 3). There was no detectable general relationship between development time *CV* and rearing temperature (Figure 4).

**Figure 3.**
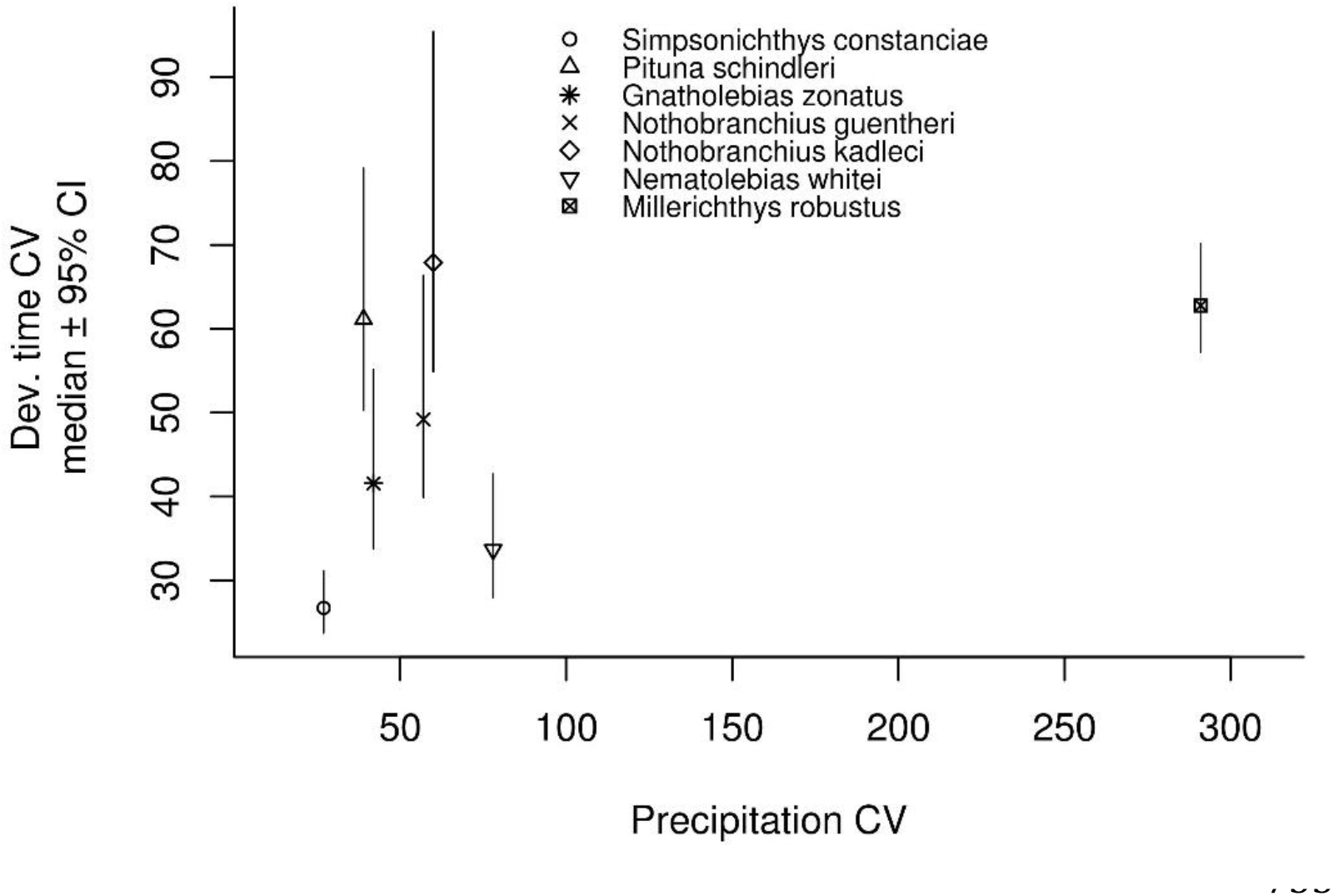
Medians of posterior distributions of species-specific coefficients of variation in development time, and their 95% credibility intervals (y-axis), against precipitation *CV* values (x-axis).

**Figure 4.**
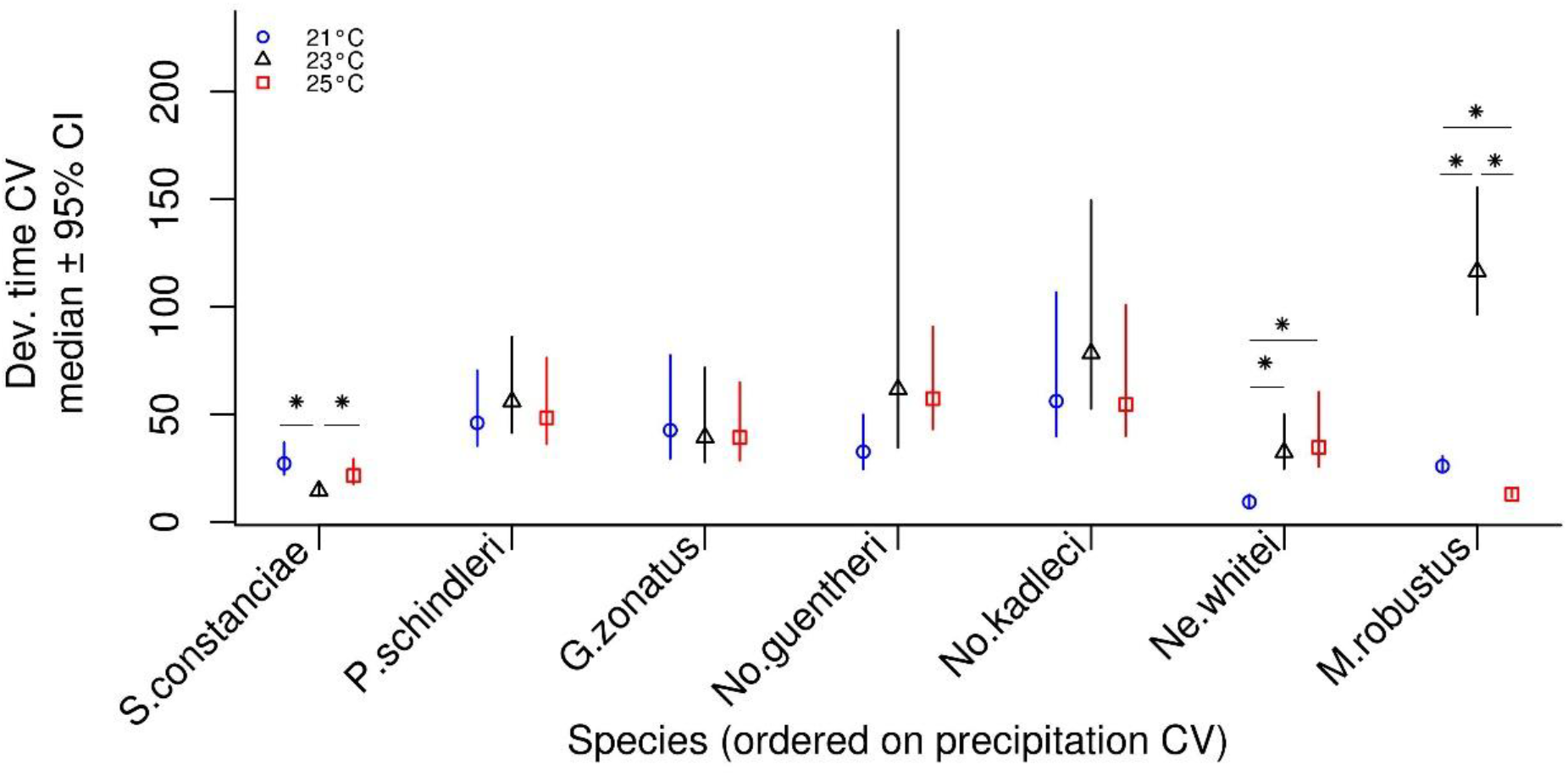
Medians of posterior distributions of species- and temperature-specific coefficients of variation in development time, and their 95% credibility intervals (y-axis). Species are ordered on a categorical x-axis scale, according to precipitation *CV* values, from the lowest (left) to the highest (right), for clarity of the results. Star symbols indicate significant within-species differences between the temperature treatment groups. A similar plot, but with continuous x-axis scale can be found in supplementary information 4, Fig. S2.

## DISCUSSION

We found substantial among-species differences in both the mean and the variation of egg development time in annual killifish species, which originate from environments along a gradient of precipitation variability. Under precipitation-driven bet-hedging scenario, we expected species from more unpredictable environments to have higher variation in embryo development times (Crean and Marshall 2009). However, we did not find any general relationship between variation in precipitation and variation in development time across species. Moreover, in several species, both the mean and the variation of development time were dependent on rearing temperature. The lack of association between variation in development time and environmental variation suggests that bet-hedging either may not be an important mechanism for persisting in these ephemeral habitats, or that other environmental factors, which the species are exposed to in the wild, also influence developmental times. Hence, if species specific variances are adaptive, the relationship between development and variation in precipitation is complex, and does not diverge in accordance with simple linear relationships.

In contrast to our results, comparative analyses have shown that annual desert plant species occurring in areas with unpredictable rates of precipitation, produce seeds with highly variable germination times (Venable 2007; see also Evans and Donnehy 2005). Diapausing eggs in the killifish system could be viewed as analogous to the example of seed-banks in plants. However, although the seasonal pond filling period, and thus precipitation, is crucial for the survival of annual killifishes (Watters 2009; Polačik et al. 2011; Domínguez-Castanedo et al. 2017), we did not find any clear relationship between precipitation predictability and variation in embryo development time. Our results are hence similar to a recent study that did not find any differences in development times among eight populations of two closely related annual *Nothobranchius* killifish species (*No. furzeri* and *No. kadleci*; Polačik et al. 2018). Together, these results suggest that other factors likely also contribute to driving differences in egg development patterns across species. While we found no evidence for bet-hedging, our results implicate a link to environmental conditions. For example*, Ne. whitei and S. constanciae* co-occur in areas with two rainy seasons per year (Costa 2012), and despite key differences in other traits between these two species (e.g. egg size and growth rate, Huber et al. 2016; Eckerström-Liedholm et al. 2017; Sowersby et al. 2019), egg development time was similar, both in terms of the variation (relatively low), mean (relatively long), and plastic to incubation temperature. These phenotypic similarities among two co-occurring species suggest that developmental time might be shaped more by the natural conditions under which these species occur.

In a bet-hedging scenario, there will be constant selection against a non-trivial part of the population that is not suited to current environmental conditions (Kussell & Leibler 2005; Beaumont et al. 2009). However, a lack of bet-hedging, particularly in unpredictable environmental conditions, may also be costly. For example, without a bet-hedging strategy, all offspring may be ill-equipped to cope under current environmental conditions. Variation in one key trait, development time, may therefore evolve as an evolutionary trade-off between the costs and benefits of employing a bet-hedging strategy. Hence, one potential explanation for our results may be that different species exhibit different solutions to this trade-off, by either trying to match the rainy season (high risk – high payoff strategy), or by having large variation (low risk – low payoff strategy) in development rates. Either scenario could yield the pattern we observed; with large among species differences in egg development rates, which were not necessarily linked to environmental predictability.

Plastic responses and adjustments in trait variance are important for facilitating adaptation, particularly as a warming climate changes both the environment and its predictability (Robeson 2002; Thornton et al. 2014; Berg and Hall 2015). Species that utilize bet-hedging strategies are hypothesized to be better adapted to cope with increased environmental variation, for instance, due to climate change, as they use a risk-minimizing strategy (Childs et al. 2010). Hence, in our study, species with higher variation in development time (for example *P. schindleri*, *No. kadleci*, and *M. robustus*), may be predisposed to cope better with future environmental changes, however at the same time they are likely to suffer significant costs, as a large proportion of a female’s eggs are unlikely to develop in any particular season.

In addition to differences among species, in variation in embryo development time, we found that both the mean and the variation in development time were affected by rearing temperature. In regard to mean development time, this result was as expected, as thermal plasticity is pronounced in ectotherms, and generally increases the speed of biological processes (Roff 2002; for killifish diapause specifically, see Levels and Denucé 1988; Podrabsky et al. 2010; Furness et al. 2015a). For three of the species in our study (*S. constanciae, P. schindleri,* and *Ne. whitei*), we found this expected pattern of typically ectotherm development, with higher rearing temperatures speeding-up developmental times. In general, phenotypic plasticity is thought to correlate with thermal variability (Stillman and Somero 2000; Angilletta 2009), as species encountering a wide range of thermal conditions should be adapted to cope with a wider range. Indeed, in two temperate-living Neotropical species (*S. constanciae, Ne. whitei*), we observed this expected negative correlation between diapause duration and ambient temperature.

We found some clear patterns between incubation temperature and mean embryo development time, although patterns of temperature-mediated changes in the variation of developmental time *(coefficient of variation)* were less clear. However, there were some significant differences among the variations estimated for each thermal incubation regime. Again, differences among the coefficients of within-species variation, across thermal regimes, tended to occur in species from cooler areas (South: *S. constanciae, Ne. whitei,* North: *M. robustus*). However, across these species there was no clear directionality in how the different temperatures affected the coefficient of variation for development time. For example, while *M. robustus* had the highest variation in the intermediate incubation temperature (23°C), *S. constanciae* had the lowest variation in this intermediate temperature. Our results hence suggest that evolution under more temperate thermal conditions both increases the evolution of thermal plasticity, and decreases the canalization of traits. However, it is difficult to make any inferences on the adaptive value of increased variations across the different temperatures.

Factors such as maternal age (Podrabsky et al. 2010), egg-laying order (Polačik et al. 2016a), and hormonal levels in mothers (Pri-Tal et al. 2011) have been reported to also influence variance in developmental rates of annual killifish embryos. We found no evidence that any maternal factors influenced developmental times in our study (maternal age and group/single breeding). We did not have sufficient statistical power to test whether females or males differed in terms of variation in the development time of the eggs/embryos they produced. However, species variance accounted for a large part of the variation in the model, with little influence of male and female effects, suggesting that species have instead evolved different development times due to differences in their natural environment. In addition to maternal influence, several other environmental factors have been found to induce plastic effects on the length of embryonic development, such as temperature (Levels and Denucé 1988; Podrabsky et al. 2010; Furness et al. 2015a), photoperiod (Levels and Denucé 1988; Podrabsky and Hand 1999; Furness et al. 2015a), hypoxia (Inglima et al. 1981), and the presence of other fish in aquaria with developing embryos (Inglima et al. 1981; Levels et al. 1986). While these factors are mostly standardized in our experiment, we cannot exclude the possibility that they affected our results. In our study, we kept developing embryos in water (solution), which allowed for highly standardized conditions, but which could decrease development times (Polačik et al. 2016b). Specifically, we cannot exclude the possibility that species differ in how they respond to developing in water (opposed to developing buried in substrate), which could in part explain our results. Further, we used captive bred fish populations, which may mean that some of the species could be relatively inbred. However, all but one species have been collected from the wild within the past 11 years, suggesting that inbreeding effects have had a relatively short time to accumulate (Table S1, supplementary information 1). Finally, as inbreeding is considered to decrease genetic variation, we note that virtually all of the species showed considerable levels of variation in development times (CV >20). Hence, we do not believe that inbreeding has driven the patterns observed in our results.

In conclusion, we found that means and variation in developmental times differed among seven annual killifish species, irrespective of environmental predictability. Moreover, we found among-and within-species differences in response to temperature treatments, with more pronounced changes in developmental time than developmental time *CV,* and with species from more temperate areas being more plastic, compared to species from more tropical areas. Although we were unable to pinpoint the exact causes of these observed differences, we suspect that a combination of environmental factors play an important, but as of yet unidentified role, in influencing embryo development times. Therefore, it is important to investigate factors potentially influencing embryo development times encountered in the natural environment of any particular species, especially with climate change further increasing environmental unpredictability.

## Acknowledgements

We thank Niclas Kolm and John Fitzpatrick for practical help, Bertil Borg for constructive comments on the manuscript, Alma Tiwe for assistance with animal husbandry, and Olivia Selander for help with experimental procedures. This research was supported by The Swedish Research Council grant to B.R. (nr. 2013-05064). The authors declare no conflicts of interests.

## Data archiving

Data will be deposited to the Dryad archive together with article publication.

