## Supplementary Information for "Variation in developmental rates is not linked to environmental unpredictability in annual killifishes"

**Supplementary information 1:** Information on laboratory killifish populations and climate data calculated based on among-years means and standard deviations retrieved from New_LocClim (FAO 2018).

Table S1. Population-specific data with population-specific and species-wide coefficients of variation (*CV*s) across their distributions, in among-year precipitation during the rainy season months.

| Species | Population, collection year | Climate data coordinates | Rainy season months | Population-specific precipitation mean (mm) | Species-wide precipitation mean (mm) | Population-specific precipitation CV (%) | Species-wide precipitation CV (%) |
|---|---|---|---|---|---|---|---|
| Gnatholebias zonatus | Finca Las Palmas, Colombia, 2014 | 4.004N:73.167W | mar, apr, may | 311 | 242 | 40,9 | 42 |
| Gnatholebias zonatus | Las Mercedes, Venezuela, 2014 | 9.10N:66.39W | may, jun, jul | 173 | | 44,3 | |
| Millerichthys robustus | Tlacotalpan, Veracruz, Mexico, 2017 | 18.627N:95.648W | jun, jul, aug | 434 | 434 | 257,9 | 258 |
| Nematolebias whitei* | Buzios, Brazil: *first rainy season* | 22.833S:41.883W | mar, apr, may | 52 | 69 | 62,8 | 78 |
| Nematolebias whitei* | Buzios, Brazil: *second rainy season* | 22.833S:41.883W | oct,nov,dec | 87 | | 92,7 | |
| Nothobranchius guentheri | Zanzibar 2014 | 5.017S:39.750E | mar, apr, may | 352 | | 89,5 | |
| Nothobranchius guentheri | Zanzibar 2014 | 6.025S:39.328E | mar, apr, may | 311 | 317 | 50 | 57 |
| Nothobranchius guentheri | Zanzibar 2014 | 6.167S:39.367E | mar, apr, may | 288 | | 33,2 | |
| Nothobranchius kadleci | Pungwe, Moçambique, 2012 | 19.291S:34.231E | dec, jan, feb | 170 | | 54 | |
| Nothobranchius kadleci | Nhamatanda, Moçambique, 2011 | 20.688S:34.107E | dec, jan, feb | 201 | 181 | 62,2 | 60 |
| Nothobranchius kadleci | Save, Gorongose, Moçambique, 2008 | 21.015S:34.463E | dec, jan, feb | 171 | | 65,5 | |
| Pituna schindleri | União, Piauí, Brazil | 4.674S:42.005W | dec, jan, feb | 203 | 203 | 39,2 | 39 |
| Simpsonichthys constanciae | Barra de Sao Joao, Brazil, 1995 | 22.030S:42.020W | oct, nov, dec | 186 | 186 | 27,3 | 27 |

*in case of N. whitei, data concerning the same population is presented in separate rows for the two rainy seasons.

**Supplementary information 2:** Information on parental generation and egg number

Table S2. Number of families, breeding males and females per species, and number of surviving eggs per family.

| family | species | female | male | egg number |
|---|---|---|---|---|
| 1 | constanciae | con1a | con1 | 12 |
| 2 | constanciae | con1a | con7 | 2 |
| 3 | constanciae | con1a | con8 | 2 |
| 4 | constanciae | con1a,2a,6a | con1 | 2 |
| 5 | constanciae | con2a | con3 | 13 |
| 6 | constanciae | con2s | con10 | 6 |
| 7 | constanciae | con2s | con3 | 3 |
| 8 | constanciae | con2s | con7 | 8 |
| 9 | constanciae | con2s,3s | con3 | 1 |
| 10 | constanciae | con3a | con10 | 3 |
| 11 | constanciae | con3a | con3 | 12 |
| 12 | constanciae | con3a | con9 | 4 |
| 13 | constanciae | con3s | con1 | 5 |
| 14 | constanciae | con3s | con8 | 3 |
| 15 | constanciae | con3s | con9 | 4 |
| 16 | constanciae | con4a | con6 | 9 |
| 17 | constanciae | con4s,5s,6s | con9 | 3 |
| 18 | constanciae | con5a | con10 | 2 |
| 19 | constanciae | con5a | con7 | 5 |
| 20 | constanciae | con6a | con1 | 1 |
| 21 | guentheri | gue4a | gue4 | 1 |
| 22 | guentheri | gue1a | gue4 | 10 |
| 23 | guentheri | gue1s | gue1 | 1 |
| 24 | guentheri | gue2.1s-2.3s | gue2.3 | 4 |
| 25 | guentheri | gue2.4-2.7 | gue2.2 | 1 |
| 26 | guentheri | gue2.4s-2.6s | gue1 | 14 |
| 27 | guentheri | gue2.4s-2.7s | gue2.2 | 1 |
| 28 | guentheri | gue3s | gue3 | 3 |
| 29 | guentheri | gue4a | gue4 | 1 |
| 30 | guentheri | gue_group1 | gue3.4 | 5 |
| 31 | guentheri | gue_group2 | gue3.1 | 1 |
| 32 | kadleci | kad2a | kad1 | 2 |
| 33 | kadleci | kad3a | kad8 | 1 |
| 34 | kadleci | kad4a | kad1 | 1 |
| 35 | kadleci | kad4a | kad4 | 3 |
| 36 | kadleci | kad4a | kad9 | 6 |
| 37 | kadleci | kad4a,6a | kad2.1 | 1 |
| 38 | kadleci | kad4s | kad7 | 6 |

Table S2 cont.

| | | | | |
|---|---|---|---|---|
| 39 | kadleci | kad5a | kad2 | 1 |
| 40 | kadleci | kad5a | kad4 | 1 |
| 41 | kadleci | kad5a | kad5 | 9 |
| 42 | kadleci | kad5s | kad11 | 6 |
| 43 | kadleci | kad6a | kad9 | 2 |
| 44 | robustus | rob1s,2s | rob1 | 187 |
| 45 | robustus | rob1s,2s | rob2 | 1 |
| 46 | robustus | rob2s | rob2 | 98 |
| 47 | robustus | rob_1 | rob1 | 3 |
| 48 | robustus | rob_group1 | rob1 | 2 |
| 49 | schindleri | schi_group1 | schi_1 | 33 |
| 50 | schindleri | schi_group2 | schi_1 | 24 |
| 51 | whitei | whit1a | whit2 | 1 |
| 52 | whitei | whit1a | whit8 | 1 |
| 53 | whitei | whit1s | whit2 | 6 |
| 54 | whitei | whit1s | whit3 | 3 |
| 55 | whitei | whit1s | whit7 | 3 |
| 56 | whitei | whit2s | whit1 | 1 |
| 57 | whitei | whit2s | whit6 | 2 |
| 58 | whitei | whit2s | whit8 | 2 |
| 59 | whitei | whit3a | whit1 | 1 |
| 60 | whitei | whit3s | whit5 | 1 |
| 61 | whitei | whit4a | whit1 | 7 |
| 62 | whitei | whit4a | whit2 | 1 |
| 63 | whitei | whit4a | whit6 | 4 |
| 64 | whitei | whit4a | whit7 | 5 |
| 65 | whitei | whit4s | whit1 | 3 |
| 66 | whitei | whit5s | whit2 | 1 |
| 67 | whitei | whit5s | whit5 | 1 |
| 68 | whitei | whit6s | whit2 | 2 |
| 69 | whitei | whit7a | whit9 | 2 |
| 70 | whitei | whit7s | whit10 | 2 |
| 71 | zonatus | zon_group1 | zon1 | 6 |
| 72 | zonatus | zon2a | zon5 | 1 |
| 73 | zonatus | zon2a,5a | zon7 | 2 |
| 74 | zonatus | zon4s,5s | zon2 | 5 |
| 75 | zonatus | zon4s,5s | zon4 | 8 |
| 76 | zonatus | zon4a6a8a | zon3 | 4 |
| 77 | zonatus | zon4a6a8a | zon7 | 1 |
| 78 | zonatus | zon4s,5s | zon2 | 8 |

**Supplementary information 3:** The models

Table S3. List of the models.

| Model nr. | Model testing | Model formula |
|---|---|---|
| 1a | Female effects, run on the data including single-female tanks only | development time ~1, random = ~ spec+male+female |
| 1b | Species and male effects, run on full data | development time ~1, random = ~ spec+male+female |
| 2 | Differences among species means and variances, pooled temperature treatments | development time ~1, random = ~ spec, rcov=~idh(spec):units |
| 3 | Differences among species means and variances, 21°C | development time ~1, random = ~ spec, rcov=~idh(spec):units |
| 4 | Differences among species means and variances, 23°C | development time ~1, random = ~ spec, rcov=~idh(spec):units |
| 5 | Differences among species means and variances, 25°C | development time ~1, random = ~ spec, rcov=~idh(spec):units |

**Supplementary information 4:** Additional result figures

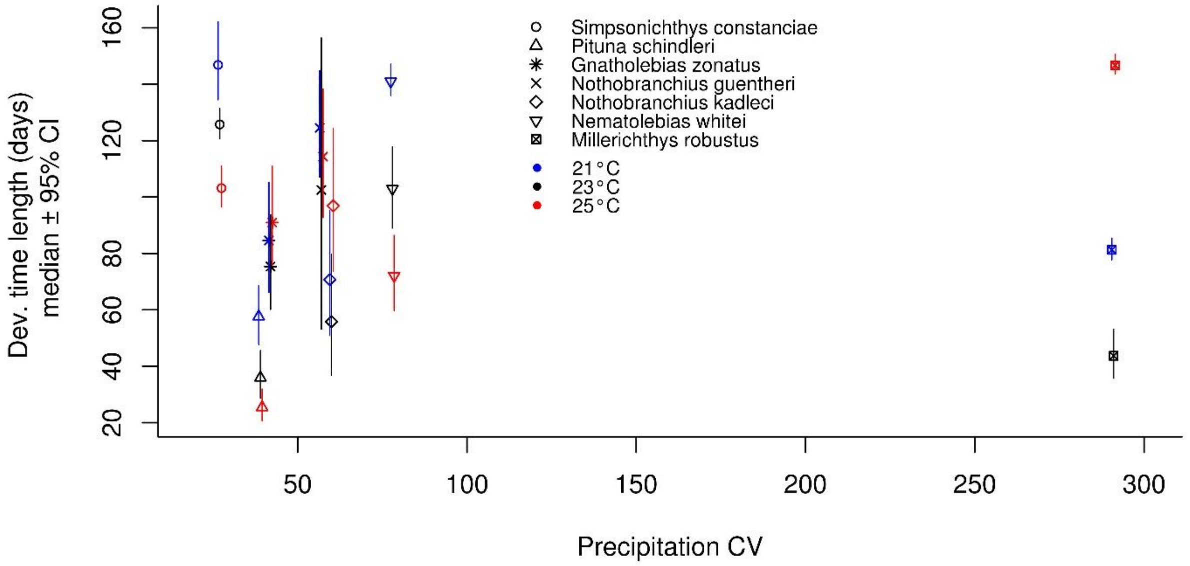


Figure S1. Species- and rearing temperature-specific medians of development time length, and their 95% credibility intervals (y-axis), against precipitation CV (x-axis).

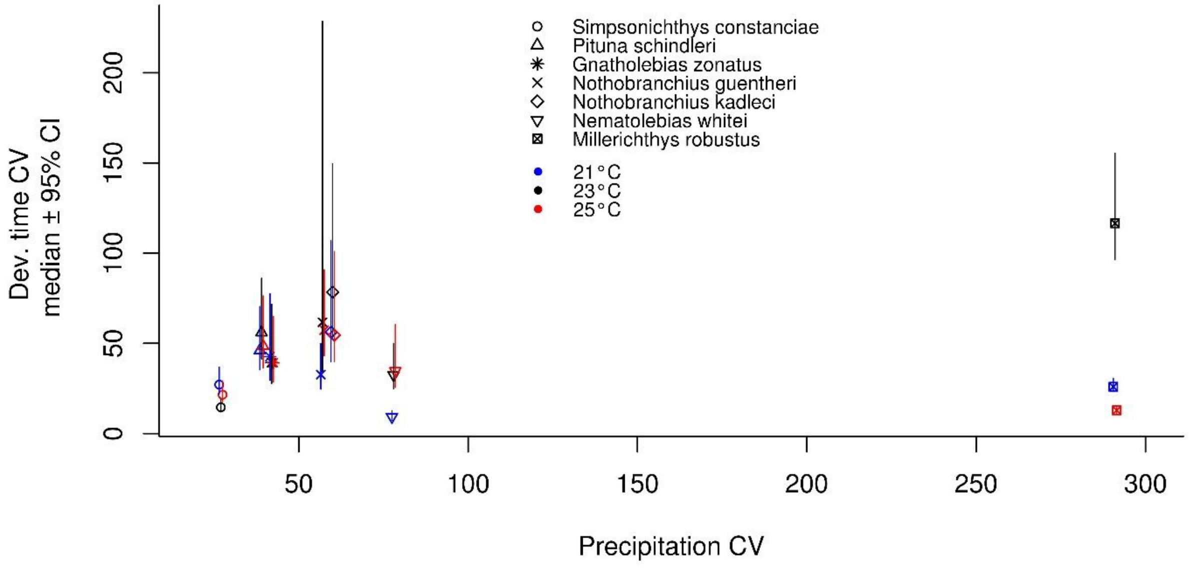


Figure S2. Species- and rearing temperature-specific medians of development time CV, and their 95% credibility intervals (y-axis), against precipitation CV (x-axis).
